# TrackUSF, a novel methodology for automated analysis of ultrasonic vocalizations, reveals modified social communication in a rat model of autism

**DOI:** 10.1101/575191

**Authors:** Shai Netser, Guy Nahardiya, Gili Weiss-Dicker, Roei Dadush, Yizhaq Goussha, Hala Harony-Nicolas, Lior Cohen, Kobi Crammer, Shlomo Wagner

## Abstract

Rodents emit various social ultrasonic vocalizations (USVs), which reflect their emotional state and mediate social interaction. USVs are usually analyzed by manual or semi-automated methodologies categorizing discrete USVs according to their structure in the frequency-time domains. This laborious analysis hinders effective use of USVs for screening animal models of human pathologies associated with modified social behavior, such as autism spectrum disorder (ASD). Here we present a novel, automated methodology for analyzing USVs, termed TrackUSF. To validate TrackUSF, we analyzed a dataset of mouse mating calls and compared the results with a manual analysis by a trained observer. We found that TrackUSF was capable of detecting most USVs, with less than 1% of false-positive detections. We then employed TrackUSF to social vocalizations in *Shank3*-deficient rats, a rat model of ASD and found, for the first time, that these vocalizations exhibit a spectrum of deviations from pro-social calls towards aggressive calls.

## Introduction

Vocal communication is fundamental to social interactions of most vertebrate species (Krams et al., 2012; McComb and Semple, 2005; Pollard and Blumstein, 2012). In humans, vocal communication is highly dynamic, with distinct vocal signals characterizing different types of social interactions and reflecting distinct emotional states (Liebenthal et al., 2016; Nygaard and Queen, 2008). In diverse social contexts and activities, such as parenting, mating, fighting and playing, mice and rats also emit various types of vocal signals, mostly at the ultrasonic range (Portfors, 2007). Such ultrasonic vocalizations (USVs) reflect the animal’s emotional state and facilitate or inhibit social interaction (Brudzynski, 2013; Knutson et al., 2002; Wohr and Schwarting, 2013). Therefore, they have gained interest as a proxy model for speech and language (Arriaga et al., 2012; Castellucci et al., 2016; Fischer and Hammerschmidt, 2011a) as well as for affective vocal communication in humans (Burgdorf et al., 2011; Panksepp, 2007).

Notably, USVs can be easily recorded and followed across extended periods of time, by simply positioning an ultrasonic microphone in the animals’ vicinity. The easiness of this approach makes the identification of modified social vocalizations in animal models of pathological conditions a promising method for monitoring changes in social behavior and for screening potential therapeutics for such conditions (Scattoni et al., 2009; Schwarting and Wohr, 2012; Wohr and Scattoni, 2013). Indeed, modified social vocalization activity was previously studied in various mouse models of autism spectrum disorder (ASD), where impairment in social communication is a core symptom (Fischer and Hammerschmidt, 2011b; Kazdoba et al., 2016; Wohr, 2014). However, the analysis of social vocalizations in animal models is usually performed by manual or by semi-automated methodologies aiming to extract discrete USVs from the audio recording and to categorize them according to their structure in a spectrogram (Brudzynski, 2009; Heckman et al., 2016; Portfors, 2007). These highly laborious and observer-dependent methodologies hinder an efficient and large-scale use of such approach for monitoring changes in social communication in animal models.

Here we present a novel methodology; the TrackUSF, and tools that we have developed to analyze ultrasonic vocal communication in rodents in an automated manner. This methodology, which avoids detection and characterization of discrete USVs, is based on a technology that is commonly used for human speech detection (see for example Arias-Londono et al., 2011; Mei et al., 2019; Nasr et al., 2018; Vergin et al., 1999). We validated the usefulness of our TrackUSF methodology by using it to analyze mouse mating calls and comparing the results to those obtained by the traditional USV-based approach. While doing this, we revealed a difference in the frequency of mating calls between two common laboratory mice trains. We then demonstrated the efficacy of our methodology in identifying modified social vocalizations in animal models, by revealing, for the first time, impaired reciprocal vocal communications in adult male *Shank3*-deficient rats.

## Results

### Validation of the novel TrackUSF methodology using mice mating calls

The TrackUSF methodology, designed to process and analyze an auditory recording in an automated manner, is schematically described in Fig. 1A. Briefly, the auditory clip is segmented into 6-msec fragments. All fragments that pass a 15 kHz high-pass filter and contain signals above a predetermined power threshold in the ultrasonic range (up to 100 kHz) are collected, and are herein termed ultrasonic fragments (USFs). The power spectrum between 15-100 kHz of these USFs is then transformed to mel-frequency cepstral coefficients (MFCCs). USFs from all audio clips of the experiment are then analyzed together using a 3-dimentional (3-D) T-distributed Stochastic Neighbor Embedding (t-SNE) tool for visual clustering of the multi-dimensional dataset (see Methods).

**Figure 1.**
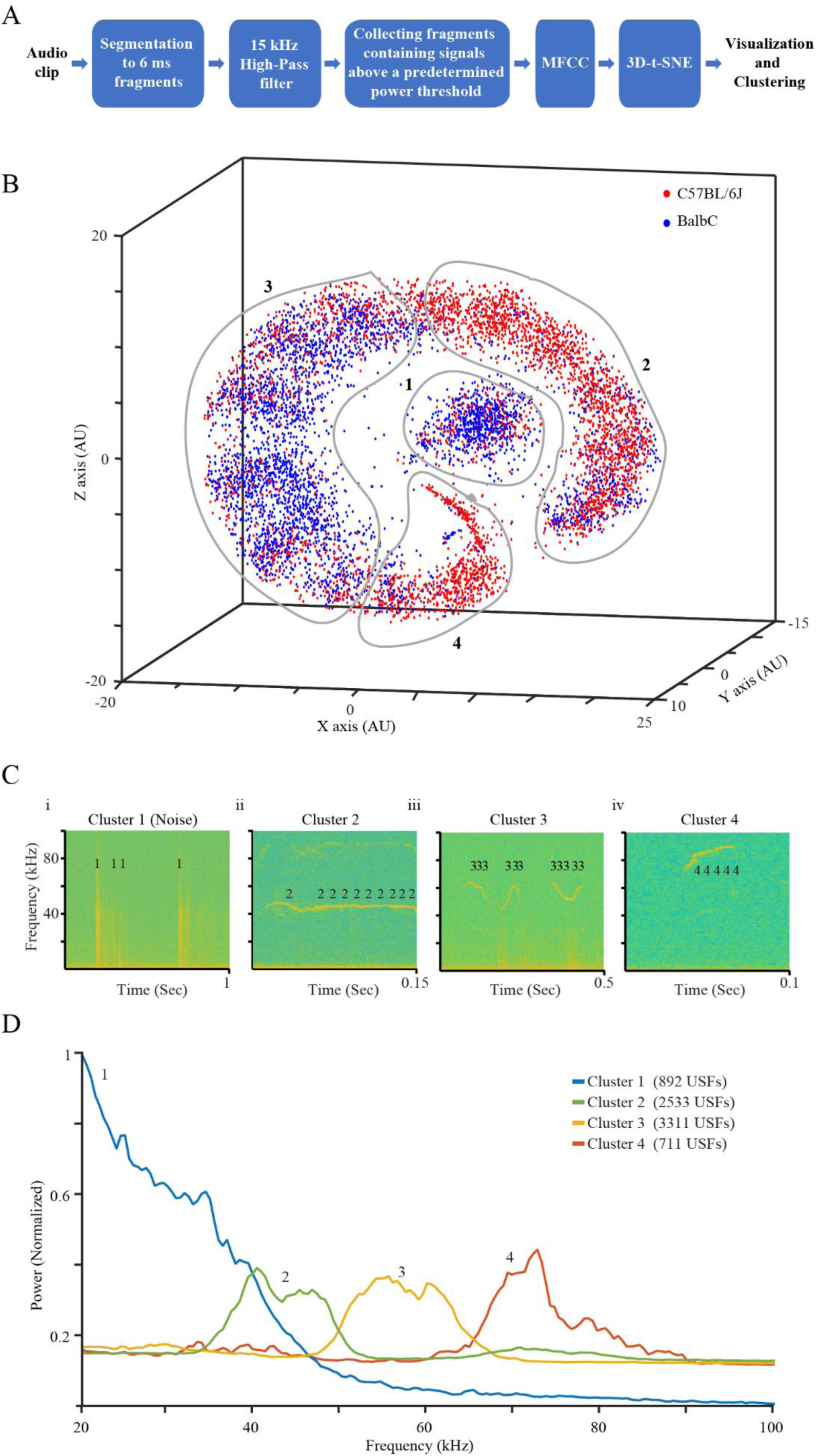
Automated analysis of mouse mating calls using TrackUSF. A) The processing pipeline used for analysis of ultrasonic vocalizations by TrackUSF. B) 3D t-SNE analysis of all USFs recorded from three C57BL/6J and three BalbC mice pairs, following MFCC transformation. Each UFS is represented by a dot, color-coded for the strain. Black numbers represent the distinct clusters, defined by the manually drawn grey lines. Note the clear separation of cluster 1, which include non-vocal signals defined as noise. C) Examples spectrograms showing USFs from all clusters, each marked as the number of the cluster it is associated with, superimposed by the TrackUSF software on their corresponding noise (i) or USVs (ii-iv) signals. D) PSD analysis of the distinct clusters shown in B. The total number of USFs in each cluster is detailed in the legend. Note the unique profile of cluster 1, which is mainly enriched with non-vocal signals (noise).

We validated our methodology by comparing the analysis of mice mating calls between TrackUSF and the traditional USV-based methodology. To that end, we audio recorded six pairs of male and female mice from two inbred strains (C56BL/6J and BalbC, three pairs per strain), which generated six 10-min long audio clips (one per pair). Notably, analysis of these audio clips using the manual USV-based methodology took a well-trained observer about 30 hours of work, while analysis of the same audio clips using TrackUSF on a standard computer took only 15 min. Moreover, the TrackUSF software allowed us to generate a 3D t-SNE representation of all USFs (Fig. 1B, each USF is represented by a single dot), which revealed an apparent separation between USFs emitted by C57BL/6J pairs (red) and those emitted by BalbC pairs (blue). It also allowed us to visually define various clusters of USFs (Fig. 1B, grey lines) and to overlay the detected USFs onto the spectrogram of each audio-clip, thus to inspect the appearance and timing of each USF in respect to its corresponding USV. This is demonstrated by the examples shown in Fig. 1C, where groups of USFs, depicted as numbers based on their cluster affiliation in Fig. 1B, represent distinct USVs. The first example (Fig 1Ci) includes only non-vocal sounds and was enriched with USFs from cluster 1, which was clearly separated from all other clusters in our t-SNE analysis (Fig. 1B), suggesting that cluster 1 is mostly composed of non-vocal sounds (herein termed noise). The other examples include USVs represented by USFs originating from clusters 2-14 (Fig 1Cii-iv).

To further analyze each of the clusters defined in Fig. 1B, we used Power Spectral Density (PSD) analysis to characterize the power spectrum of USFs from each of the clusters, while focusing our analysis the ultrasonic range between 20-100 kHz. As apparent in Fig. 1D, all clusters, except for cluster 1, showed clear distinct peaks at specific frequencies. In contrast, cluster 1 included USFs of variable frequencies, mostly at the lower range. Given this and our findings suggesting that these USFs represent noise, cluster 1 was excluded from all downstream analyses.

To directly compare the results obtained using TrackUSF to those achieved with the manual USVs-based methodology, we plotted the distribution of both, the TrackUSF-detected USFs for each cluster and the manually detected USVs over time. As exemplified in Fig 2A, USVs (manually detected) appeared in sequences, with prolonged periods of silence between them. Notably, almost all USVs were represented by at least one USF (from cluster 2-5), with no false positive USFs. To examine the influence of the power threshold used for TrackUSF on the overlap between USVs and USFs, we employed TrackUSF to analyze the data using five distinct threshold levels (1, 1.5, 2.2, 2.7 and 3.5 [arbitrary units]). It is noteworthy to mention that the time it took our software to analyze the data set ranged between 10 min for the highest and 120 min for the lowest threshold. The percent of manually detected USVs which were represented by at least one USF ranged between 84% in the lowest threshold (threshold=1) to 46% in the highest threshold (threshold=3.5) (Fig. 2B). The total duration of manually defined USVs that was also covered by USFs ranged between 48% of time in the lowest threshold to 19% in the highest (Fig. 2C). Among the six audio clips and for all threshold levels there was a statistically significant correlation between the number of USVs detected manually and the number of USFs detected automatically by TrackUSF (R^2^=0.81, 0.88, 0.91, 0.92, 0.93 respectively, p<0.001 for all, Fig. 2D). Regardless of threshold level, we found that only very few USFs (<1% of all USFs) were false positive (not representing any real USV, Fig. 2E). Thus, for threshold of 2.7 and below, Track USFs was able to detect most manually extracted USVs with very high accuracy.

**Figure 2.**
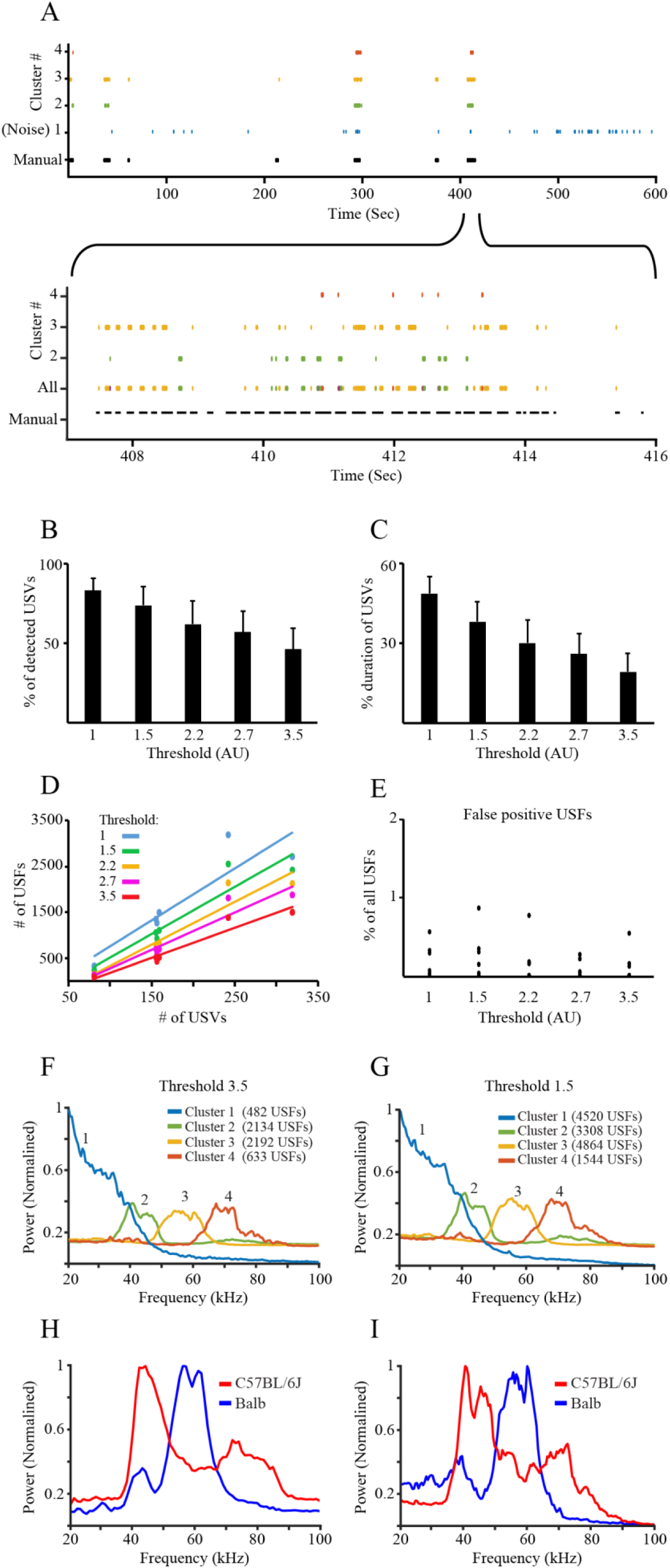
TrackUSF accurately captures most of the manually detected USVs and enables their further characterization. A) Above: co-localization of USFs (colored dots) and USVs (black dots) during a whole 10-min long audio recording of mouse mating calls. Note the various sequences of USVs, separated by prolonged silent periods. Below – One USV sequence displayed in higher resolution, with the co-localized USFs (excluding cluster 1). Note the accurate detection of most USVs by the various types of USFs. B) Percent of manually defined USVs that are detected by at least one USF, using different thresholds (1, 1.5, 2.2, 2.7, 3.5) for TrackUSF for analyzing the same mating calls dataset. C) Percent coverage of the total duration of manually defined USVs by the various USFs, for the various thresholds. D) Number of detected USFs plotted as a function of number of manually detected USVs, for the various thresholds. E) Percent of all detected USFs from all clusters, except for cluster 1, that were found to represent non-USV fragments (false-positive USFs), using different thresholds. Each point represent a distinct audio clip. Note that in none of the audio clips we observed >1% false positive USFs, regardless of threshold level. F) PSD analysis of the various clusters using a very high threshold of 3.5, as compare to 2.7 in Fig. 1D. G) As in F, for a low threshold of 1.5. Note that the PSD profile seems to be insensitive to the threshold applied. H) PSD analysis of the manually detected USVs analyzed separately for USVs emitted by C57BL/6J (red) and BalbC (blue) mice pairs, showing a tendency of BalbC mice for higher-pitch mating calls. I) As is H, for the distinct clusters of USFs detected by TrackUSF using a threshold of 2.7. Note the very high similarity with H, showing that TrackUSF properly characterize the main frequency of the USVs underlying the detected USFs.

We then compared the PSD analysis of the various USF clusters for two additional threshold (1.5 and 3.5). As presented in Fig. 2F-G, PSD analyses using a either of these thresholds generated a very similar output to that generated using a threshold of (Fig. 1D). Taken together, we conclude that PSD characterization of the USFs is relatively insensitive to the threshold applied.

Finally, as noted above and as presented in Fig. 1B, USFs generated from recordings of C57BL/6J pairs showed a distribution on the t-SNE analysis that was distinct from the distribution of USFs of BalbC pairs. Therefore, we next compared the PSD analyses of the manually extracted USVs between these two strains, (Fig. 2H). Interestingly, this analysis demonstrated a clear difference between the two strains, with the USVs of C57BL/6J mice showing tendency towards lower pitch (40 kHz) as compared to the higher pitch (60 kHz) of the BalbC USVs. To verify that similar tendencies are also seen using the TrackUSF methodology (using a threshold of 2.7), we scaled the PSD curve of each cluster to the number of USFs in this cluster and then summed the curves of the scaled clusters, separately for C57BL/6J and BalbC mice. This analysis yielded PSD curves (Fig. 2I) that were highly similar to those achieved using the manually extracted USVs (Fig. 2H), also demonstrating a clear differences between the two strains thus further validating the reliability of the TrackUSF methodology.

Overall, the TrackUSF methodology enabled automated and time efficient analysis of mating calls in mice in a manner that accurately capture most emitted USVs. Moreover, the number of USFs identified using this methodology correlated very well with the number of USVs detected by the traditional USV-based analysis and the spectral characterization of calls were shown to be very similar between both methodologies.

### TrackUSF reveals modified vocalizations during social interactions in *Shank3*-deficient rats

Following the validation of the TrackUSF methodology using mice mating calls, we examined the ability of this methodology to reveal modified vocalization activity during social interactions in *Shank3*-deficient rats, a novel rat model of ASD (Harony-Nicolas et al., 2017). During male-male social interactions, adult rats emit relatively high rate of variable USVs, generally divided into two categories: (1) the “22 kHz alarm calls”, which are associated with negative states and aversive situations and are characterized by low pitch (20-30 kHz) and prolonged durations (150-3000 ms) and (2) the pro-social “50 kHz play calls”, which are further divided into flat and highly modulated (trills) USVs and are associated with positive states and appetitive situations and are characterized by high pitch (40-80 kHz) and short durations (10-150 ms) (Brudzynski, 2009; Knutson et al., 2002; Portfors, 2007; Wohr et al., 2017). To record such USVs, we conducted experiments comprised of 10-min long encounters between dyads of adult male rats of the same genotype. The encounters were simultaneously video-and audio-recorded by a video camera and an ultrasonic microphone, respectively, located both above the arena (Supplemental Fig. 2). About half of the experiments comprised encounters between unfamiliar (novel) animals and the other half between familiar animals (cage-mates). Besides the three genotypes of *Shank3*-deficient rats (wild-type (WT), heterozygous (Het) and homozygous (KO)), we performed similar experiments with age-matched adult male Sprague Dawley (SD) rats. Overall, we recorded 109 experimental sessions. The number of experiments conducted with each genotype in each familiarity level (cagemate/novel) is detailed in Fig. 3A. Note that the relatively larger number of Het sessions reflects their abundance in the litters.

**Figure 3:**
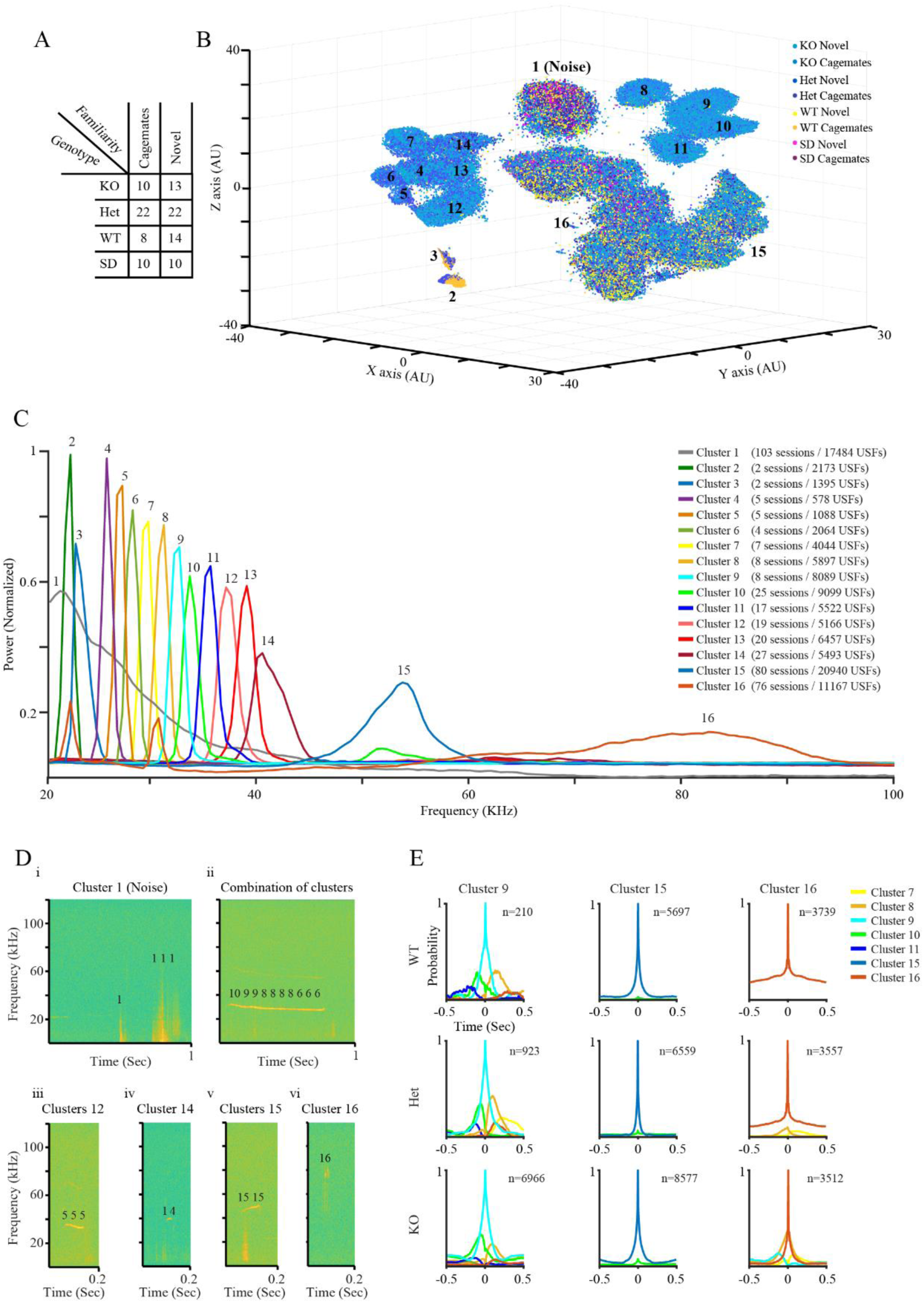
Modified pitch of ultrasonic vocalization activity revealed by our novel methodology in *Shank3*-Het and KO. A) A table that details the various types of social encounters and the number of sessions analyzed for each encounter type. B) 3D t-SNE analysis of all USFs recorded during all sessions, following MFCC transformation. Each UFS is represented by a dot, color-coded for the genotype and familiarity level. Black numbers represent the distinct clusters. Note the clear separation of cluster 1, which included non-vocal signals defined as noise. C) PSD analysis of all distinct clusters shown in B. The number of sessions represented by >10 USFs in each cluster, as well as the total number of USFs in each cluster, are detailed in the figure legend. Note the continuous spectrum in the 25-45 kHz range created by clusters 4-14. D) Examples spectrograms showing USFs from several clusters, each marked as the number of the cluster it is associated with, superimposed by the TrackUSF software on their corresponding noise (i) or USVs (ii-vi) signals. Note the trill-like appearance of USFs from cluster 16. E) Examples of vicinity curves describing the probability of a USF from any cluster (color coded for the distinct clusters) to appear before or after USF from a given cluster, across the three genotypes of *Shank3*-deficient rats. Note the stability across genotypes exemplified for clusters 9 (left) and 15 (middle), in contrast to the growing tendency of other clusters to combine with cluster 16 in Het and mainly KO animals.

All audio clips were pooled and analyzed together by TrackUSF using a threshold of As apparent in Fig. 3B, where each USF is represented by a single dot, color-coded for genotype/familiarity level, some clusters of USFs (e.g. 15, 16) included significant representation of all types of experimental sessions (all genotypes and both familiarity levels). Nonetheless, other clusters (e.g. 4-14) included almost solely USFs of Het or KO rats. These results suggested a distinct type of USVs emitted during social encounters between *Shank3*-deficient rats and their WT littermates or SD rats. To further examine this possibility, we separately analyzed the USFs represented in each cluster by PSD analysis (Fig. 3C). Similar to our analysis of mice mating calls (Fig. 1), cluster 1, which was clearly separated from all other clusters and included data from all genotypes, comprised USFs of variable frequencies at the lower range. By examining their appearance in the spectrograms, USFs of cluster 1 were found again (like in mice mating calls) to be non-vocal sounds (noise, see example in Fig. 3Di), and therefore this cluster was excluded from all further analyses. Clusters 2 and 3 were also excluded from this and other downstream analysis because they included USFs originated from only two experimental sessions (the number of experimental session representing each cluster are detailed in Fig. 3C). Clusters 4-14, which contained mainly USFs from Het and KO rats, displayed relatively sharp, well-defined peaks between 20-40 kHz. By their appearance in the spectrograms, USFs of these clusters seem to represent variations on the so-called “22 kHz alarm calls” (Fig. 3Dii-iv), thought to be associated with aggression and alarm (Brudzynski, 2013; Knutson et al., 2002; Portfors, 2007; Wohr and Schwarting, 2013). In contrast, clusters 15 and 16, which showed much wider PSD peaks ranging between 50-90 kHz (Fig. 3C), were found by us to represent the so called “50 kHz play calls” (Fig. 3Dv-vi), which as noted above, were previously associated with pro-social affiliative behavior (Brudzynski, 2013; Knutson et al., 2002; Portfors, 2007; Wohr and Schwarting, 2013). In order to examine if USFs from all other clusters (4-16) tend to appear in certain combinations, we calculated their likelihood to appear before or after a USF from a given cluster (hereafter termed vicinity), within a time window of 0.5 sec for each direction (Fig. 3E, Supplemental Fig. 3, color coded for each cluster). We found that USFs from clusters 4-14 had variable tendencies to appear in certain combinations (see for example Fig. 3E – left panels, for cluster 9), but the highest likelihood was for repetitive appearance of USFs from the same cluster, as reflected by the high amplitude of their own vicinity peak (middle peak in each representative graph in Fig. 3E and Supplemental Fig. 3). This likelihood is called by us repeatability and is further explored below. USFs of cluster 15 showed low vicinity with USFs from other clusters in all genotypes (Fig. 3E – middle panels).

Interestingly, while cluster 16 in WT rats showed the same pattern as of cluster 15, in Het and even more in KO animals this cluster showed moderate vicinity with USFs from other clusters (Fig. 3E – right panels), suggesting a deviation from pure pro-social calls in Het and KO animals, which will be further explored below. It is important to note, however, that no clear aggressive behavior was spotted by observers that examined all video clips.

### *Shank3*-deficient rats emit higher numbers of low-pitch vocalization fragments

We next examined if the number of detected USFs ranged between the various genotypes (Fig. 4A). Our analysis demonstrated that while all SD rats displayed <200 USFs, a tri-modal distribution was found in the *Shank3*-Het, KO and WT littermates, with many of them displaying >200 USFs. Nevertheless, it became clear that there were only few sessions of WT rats with >400 USFs, while most KO sessions had >600 USFs. To further explore this tendency, we categorized each session of the three genotypes of *Shank3*-deficient rats according to the number of detected USFs to low (<600) and high (>600) and examined the proportions of each genotype in these categories separately for cagemates and novel sessions. As apparent in Fig. 4B, while WT and Het animals showed a rather similar proportion of 14-27% sessions with >600 USFs, in KO animals more than 50% of the sessions were with >600 USFs. This tendency was apparent in all sessions, regardless of the familiarity between the animals (novel animals or cagemates). Statistical analysis revealed a significant difference between the three genotypes (Chi-square test, χ^2^ (5)=14.874, p=0.0109), with no familiarity-dependent differences. We therefore combined the two familiarity levels and analyzed the statistical differences between the three genotypes. We found a statistically significant difference between the three genotypes (Chi-square test, χ^2^ (2)=12.412, p=0.002). A *post hoc* analysis revealed a significant difference between KO animals and the two other groups (KO:Het - p=0.003; KO:WT - p=0.019), with no difference between WT and Het animals. We further examined this tendency separately for each of the clusters using a slightly more detailed categorization. As shown in Fig. 5A for every second cluster (see Supplemental. Fig. 4 for all 4-14 clusters), quantities of USFs from clusters 4-14, which seemed to represent the 22 kHz-like USVs, were differentially distributed between the genotypes. While WT animals showed very restricted numbers of sessions with >50 USFs, KO animals displayed high numbers of such sessions and Het animals were in between WT and KO animals. In contrast, clusters 15 and 16, which seemed to represent the 50 kHz-like USVs, were similarly distributed between the three distinct genotypes (Supplemental Fig. 5A). Thus, *Shank3*-deficient rats emit more 22 kHz-like USVs as compared to their WT littermates.

**Figure 4:**
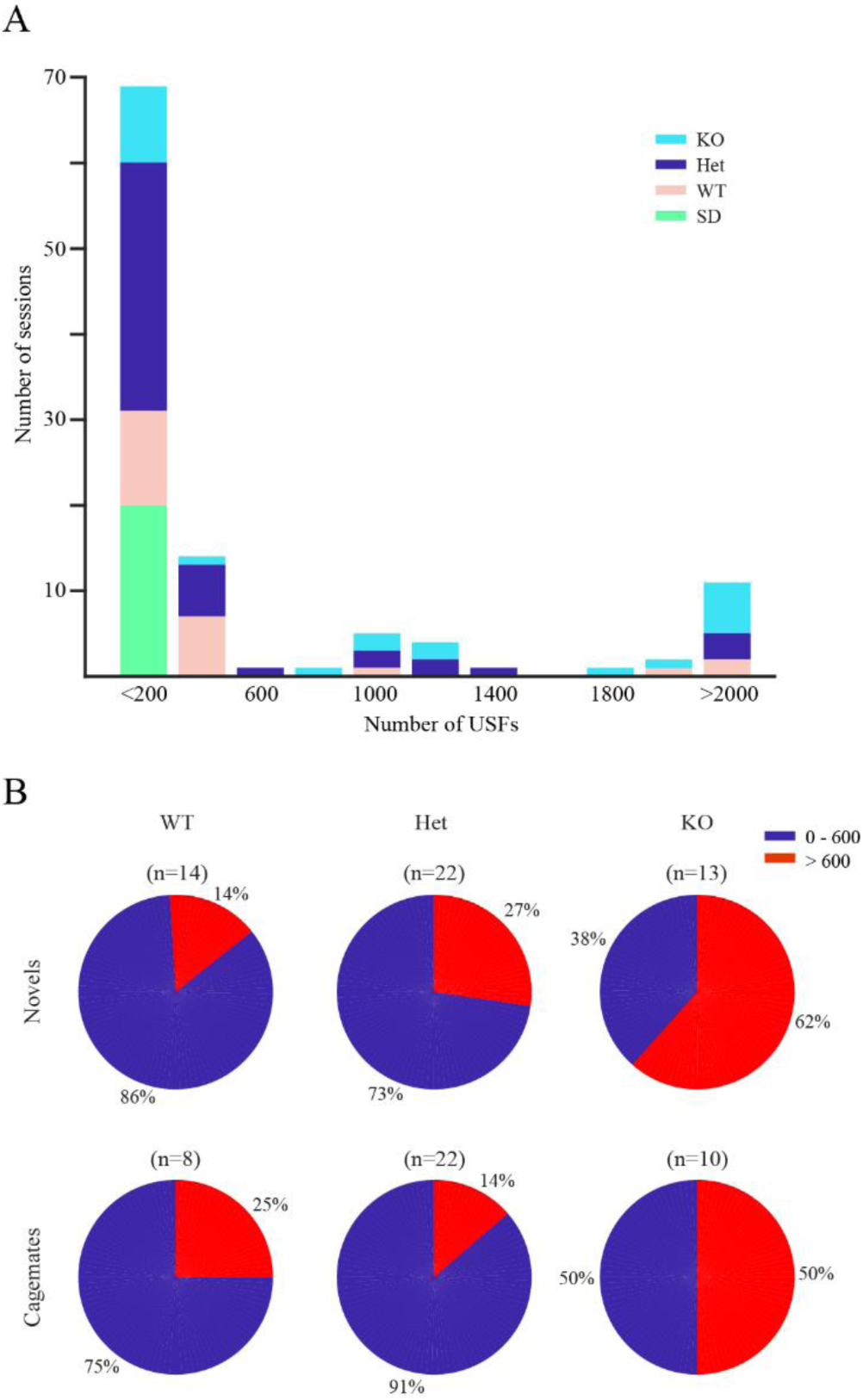
Higher numbers of USFs detected for *Shank3*-Het and KO rats, as compared to WT littermates and SD rats. A) Distributions of the sessions of each of the four genotypes examined based on the total number of USFs (excluding noise) that was detected in each session. B) Proportions of sessions with more (red) or less (blue) than 600 USFs of all clusters for the *Shank3*-Het and KO rats and their WT littermates, analyzed separately for dyads ofnovel animals and cagemates.

**Figure 5:**
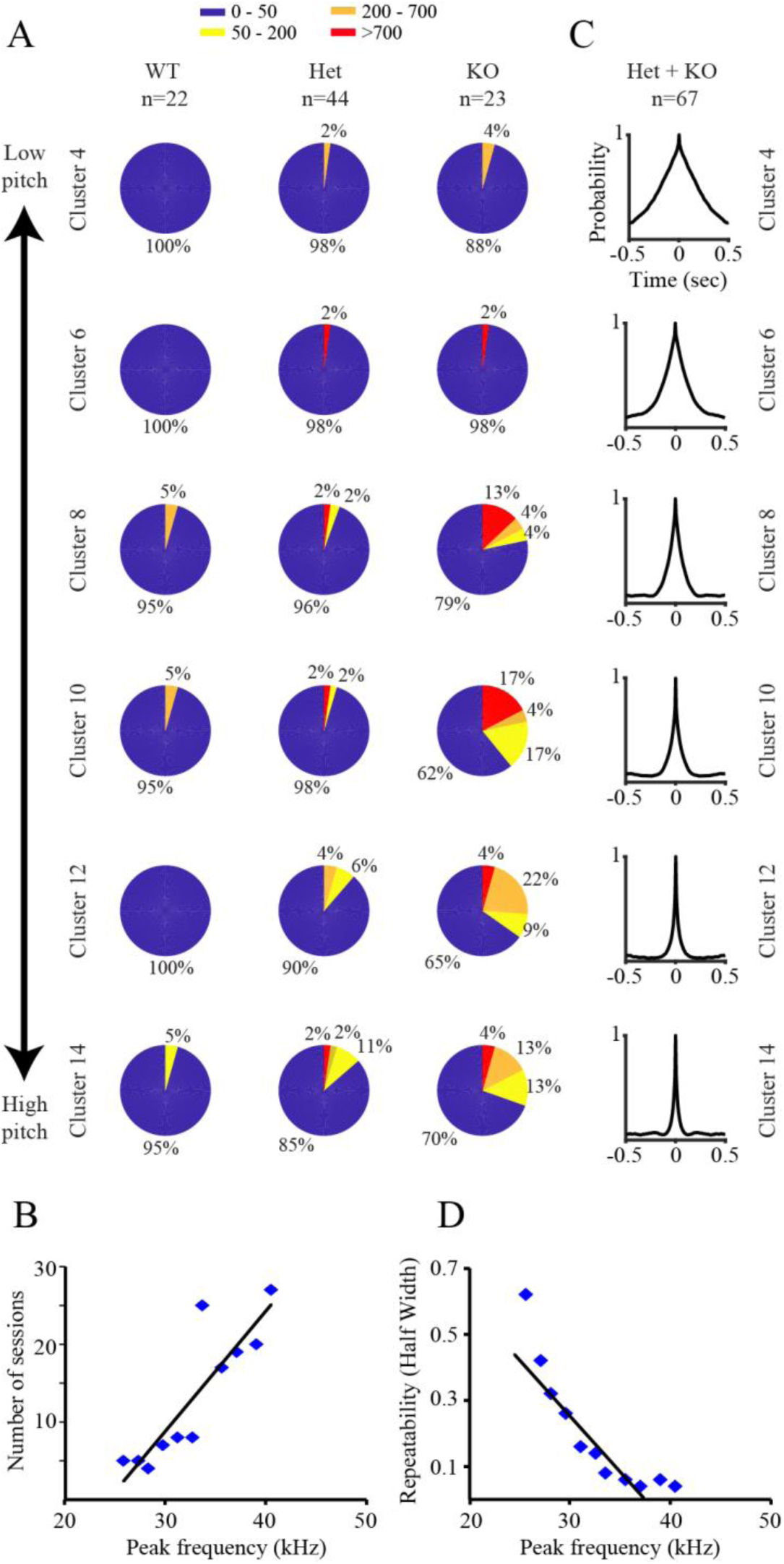
The abundance and repeatability of USFs associated with *Shank3*-Het and KO rats, correlates with their pitch. A) Proportions of sessions according to the numbers of USFs of every second cluster of clusters 4-14 for *Shank3*-Het and KO rats and their WT littermates (combining sessions of novel animals and cagemates). Note the gradual increase in proportion of sessions with high numbers of USFs, specifically exhibited by Het and KO animals. B) A statistically significant positive correlation was found between the number of sessions and PSD peak frequency of all the 4-14 clusters for *Shank3*-Het and KO rats, combined. C) Repeatability curves of each of the clusters shown in A, combined for *Shank3-*Het and KO animals. Note the gradual decrease in curve width with cluster number. D) A statistically significant negative correlation was found between the half-width of the repeatability curve and PSD peak frequency of all the 4-14 clusters for Het and KO animals, combined.

A closer look into these results suggested that there is a gradient in the number of sessions contributing for more than 50 USFs in the various clusters, with generally higher numbers of sessions for clusters representing high-pitch calls (Fig. 5A). In agreement with this observation, our statistical analysis revealed a statistically significant positive correlation (Pearson correlation, R^2^=0.77, p<0.0001) between the number of sessions contributing >10 USFs for a given cluster and the PSD peak frequency of this cluster (Fig. 5B), suggesting that high-pitch calls are more common among the various sessions. We also noticed a similar gradient, when calculating the probability of USFs to follow or precede other USFs of the same cluster (repeatability) (Fig. 5C, Supplemental. Fig. 5B for clusters 15,16). We therefore calculated the half-width of this repeatability curve for each cluster, and used it as a proxy for the duration of USVs composed of repeated appearances of the same USF. This analysis was done for Het and KO animals together, as they showed very similar repeatability curves (Supplemental. Fig. 4B), while for WT animals we did not have enough calls to perform such analysis for all clusters. A statistically significant negative correlation (Pearson correlation, R^2^=0.78, p<0.0001) was revealed between the PSD peak frequency and repeatability half-width of each cluster (Fig. 5D), suggesting that longer USVs underlie low-pitch USFs. Taken together, these results suggest that *Shank3*-deficient animals (Het and KO animals) exhibit spectrum of a behavioral impairment expressed by their modified social vocalization. Within this spectrum, a stronger impairment, exhibited by fewer animals is reflected by calls closer to 22 kHz USVs in both their pitch (low) and duration (high), while weaker and more common levels of impairment are reflected by USVs that are closer to 50 kHz calls in both pitch and duration.

### The modified vocalizations of *Shank3*-deficient rats are associated with social interactions

Finally, in order to assess whether the modified vocalization activity of *Shank3*-deficient rats is related to interaction between the animals, we examined, separately for each genotype, the distribution of USFs between periods of interaction and periods of no interaction. We first analyzed the video data using automated tracking system to identify events of physical interaction within each dyad (Fig. 6A and Methods). We then divided the interaction time to 50 msec bins and examined for each time bin whether it did or did not contain any USFs. As shown in Fig. 6B left panel, we found a statistically significant higher proportion of interaction time associated with USFs in KO animals, as compared to both WT and Het animals (Kruskall-Wallis test: χ^2^ (2)=6.713, p=0.022). No such difference was found when the same analysis was performed for time of no interaction between the animals (Fig. 6B right panel; χ^2^ (2)=4.912, p=0.0857). When this analysis was separately done for each cluster (Fig. 6C), it became apparent that this is mainly due to clusters 4-14, representing the 22 kHz-like USVs. Taken together, our data suggest that compared to their Het and WT littermates, *Shank3*-KO animals have an especially high tendency to emit 22 kHz-like USVs while interacting with each other. As these calls are associated with aggressive or defensive behaviors (Brudzynski, 2013; Knutson et al., 2002; Portfors, 2007; Wohr and Schwarting, 2013), this may suggest that as compared to interactions between their WT and Het littermates, *Shank3*-KO rats are less affiliative.

**Figure 6:**
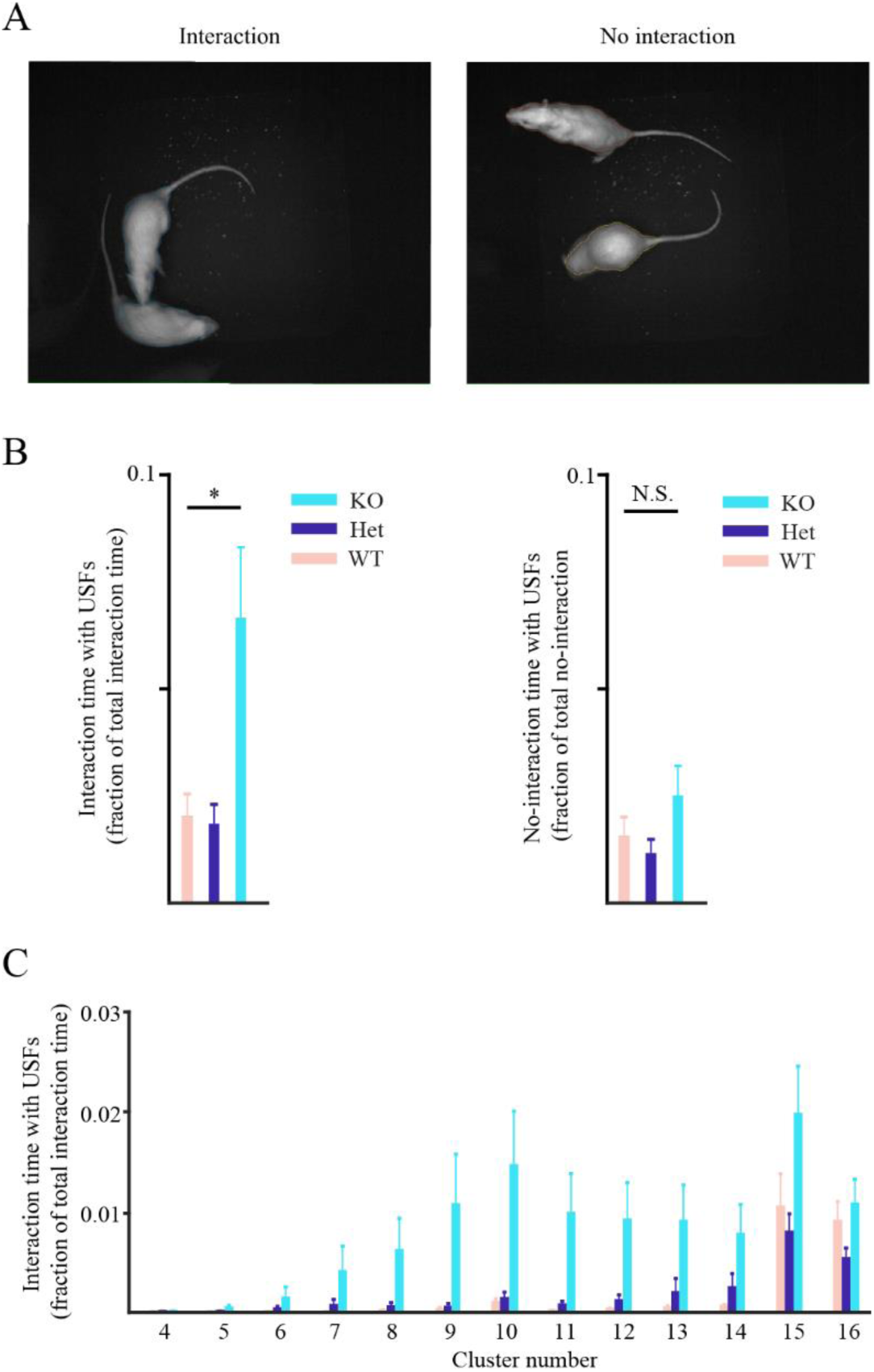
*Shank3*-KO rats show higher tendency for modified ultrasonic vocalization during social interaction. A) Representative pictures of animals during interaction (left) or no interaction (right) states, as captured by the video camera. Note the software-generated body contours used for the behavioral analysis. B) Left: The fraction of social interactions time that involves ultrasonic vocalizations (analyzed in 50 ms bins), averaged across all session separately for each genotype (combining sessions of novel animals and cagemates). Right: Same as left, for time of no social interaction. C) As in A, separately analyzed for each cluster.

## Discussion

Mice and rats are commonly used to model human pathological conditions. Under specific social contexts, these rodents emit various kinds of vocalizations, mostly within the ultrasonic range, which reflect the animal’s affective state and modulate social interactions (reviewed in Portfors, 2007). Given the relative simplicity of audio recording from animal cages, modified social vocalizations of rats and mice may serve as an excellent readout of changes in social behavior or emotional state following various manipulations (Fischer and Hammerschmidt, 2011b; Schwarting and Wohr, 2012; Wohr and Scattoni, 2013). Yet, the efficiency of such approach is limited, mainly due to the laborious methodologies used for analysis of rodents’ ultrasonic vocalizations. (Brudzynski, 2009; Heckman et al., 2016; Portfors, 2007). Recently, several computerized tools for automated or semi-automated detection and categorization of USVs were reported (Barker et al., 2014; Burkett et al., 2015; Grimsley et al., 2013; Reno et al., 2013; Van Segbroeck et al., 2017). Nevertheless, these methodologies are focused on deciphering the specific behavioral information encoded by the highly variable structures of USVs, the repertoire of which seem to get wider and wider with more and more studies performed (Burkett et al., 2015; Holy and Guo, 2005; Scattoni et al., 2011; Van Segbroeck et al., 2017; Wright et al., 2010). Yet, USV-focused analysis, as performed by current methodologies, is not suited for identifying social behavioral changes associated with pathological conditions or therapeutic interventions. To be able to identify such changes, there is a need for an efficient and automated methodology to detect differences in social vocalizations between experimental and control groups of animals. Here we developed a novel approach to explore changes in vocalization activity between groups of animals, by avoiding a direct identification of discrete USVs and rather analyzing the signature of vocalization activity in an automated manner. To this end, we adopted a methodology commonly used for human speech detection (Arias-Londono et al., 2011; Mei et al., 2019; Nasr et al., 2018; Vergin et al., 1999) and embedded it in our TrackUSF software. This methodology is based upon breaking the audio recording to short (6 msec) fragments, identifying ultrasonic fragments (USFs), transforming them using MFCC and clustering them via t-SNE analysis. The graphical user interface (GUI) of our open-source software enables loading a large number of audio clips for analysis. This feature is crucial for detecting changes between groups of animals, as the t-SNE analysis requires the inclusion all the examined audio clips at once to allow clustering relative to the entire ensemble of detected USFs. Following the t-SNE analysis, our software allows an easy extraction of specific clusters for further analysis as well as examination of USFs from any combination of clusters by their overlay on the spectrogram of a given audio clip.

We tested the efficiency of TrackUSF by analyzing two distinct sets of vocalization types, mice mating calls and rats male-male interaction calls, recorded using two different ultrasonic microphones. In both cases (Fig. 1B and Fig. 3B), we found that USFs representing non-vocal noise appeared in a distinct cluster, separated from all other USFs, which enabled their automated exclusion from further analysis. The ability of TrackUSF to automatically separate noise from all other types of vocalizations is a substantial advantage over previous methods, which spares the tedious manual steps of de-noising that is critical for analyzing these type of datasets. Furthermore, we found that USFs of other clusters, aside from the noise cluster, represent genuine USVs of various structures, with distinct clusters representing mainly USFs of different frequencies (Fig. 1C and Fig. 3D).

To further validate our methodology, we directly compared the analysis of mice mating calls using TrackUSF to the analysis of the same audio clips using the traditional methodology of manual USVs extraction by a trained observer. We found that even with the highest threshold used by us (threshold=3.5), TrackUSF was able to detect about 50% of the manually detected USVs and that this proportion rose to above 80% with lowering the threshold to 1 (Fig. 2C), with no change in the false positive detection rate (<1% of all detected USFs; Fig. 2E). The relatively low proportion (<50%) of total USV duration which is covered by USFs may be due to the fact that many USVs are broken by short periods of silence, which are not covered by USFs (see Fig. 1Ciii for an example). We also found a high correlation between the numbers of USFs and those of manually detected USVs, suggesting that changes in the number of USFs can reliably reflect changes in the number of the USVs underlying them (Fig 2D). Finally, by analyzing the mean frequency for each cluster and combining it for all clusters, we could detect the same differences in main USV frequency between C57Bl/6J and BalbC mice, as identified by analyzing manually detected USVs (Fig. 2H-I). These results confirm the efficiency of TrackUSF not only for detection of changes in ultrasonic vocalization activity but also for characterization of these changes.

To demonstrate the efficiency of TrackUSF for detecting modified social vocalizations in animal models of pathological conditions, we used it to study possible modified social vocalizations during male-male social interactions in a recently presented rat model of ASD, the *Shank3*-deficient rat (Harony-Nicolas et al., 2017). These rats were previously reported to exhibit impaired social approach behavior following playback of pro-social ultrasonic vocalization (Berg et al., 2018). The TrackUSF analysis revealed a significant number of clusters (4-14) that were mostly enriched with USFs generated by Het and KO animals (Fig. 3B) and displayed sharp peaks ranging between 25-45 kHz in the PSD analysis (Fig. 3C). In contrast, clusters 15 and 16 that included similar numbers of USFs from all genotypes, created wider peaks ranging between 50-90 kHz. This frequency range and the unique trill-like appearance of USFs from cluster 16 align well with the characteristics of the 50 kHz pro-social USVs, associated with social approach and reward (Brudzynski and Pniak, 2002; Burgdorf et al., 2001; Willuhn et al., 2014; Wohr and Schwarting, 2007, 2013). Yet, clusters 4-14 could not be simply identified as 22 kHz alarm calls, associated with aggressive behavior (Burgdorf et al., 2008; Saito et al., 2016; Schwarting and Wohr, 2012; Takeuchi and Kawashima, 1986), as their peaks created a rather continuous spectrum between 25-45 kHz (Fig. 3C).

We therefore further analyzed the duration of the USVs underlying clusters 4-14. To that end, we used the ability of TrackUSF to inform about the time point of each USF and calculated the tendency of USFs from each cluster to appear in a repeated manner, termed by us repeatedly (Fig. 5C). We then examined the relationships between the PSD peak frequency of each cluster and the half-width of its repeatability curve, which served us as a proxy for the duration of the underlying USV and found a negative correlation for clusters 4-14 |(Fig. 5D). Accordingly, USVs associated with clusters with low-frequency peaks tended to be extended, thus resembling the prolonged 22 kHz calls, while USVs associated with clusters with high-frequency peak were relatively short, similarly to 50 kHz calls (Fig. 3D). Opposite relationship was found between the frequency of the cluster PSD peak and the number of sessions contributing USFs to it; the higher the PSD peak frequency of a given cluster, the more sessions it comprised (Fig. 5B). Moreover, by counting the number of the distinct types of USFs for the distinct session we revealed that Het, and especially KO animals, emit many more USFs from clusters 4-14, than their WT littermates (Fig. 5A). Thus, during male-male interactions, *Shank3*-deficient rats seem to make enhanced use in a spectrum of vocalizations, leaning towards lower frequencies and longer durations (22 kHz-like calls). Such calls are rare in WT animals and absent from vocalizations made by SD rats, which make almost only 50 kHz pro-social call in similar experimental conditions.

To summarize, we presented here a novel methodology and open-source computerized tool, termed TrackUSF that enables automated analysis of ultrasonic vocalization activity of rats and mice. This methodology, which avoids analyzing discrete USVs, is mainly suited for detecting differences in vocalization activity between groups of animals. We validated this methodology by analyzing mice mating calls and showing that TrackUSF is capable of identifying most manually detected USVs and characterizing differences between C57BL/6J and BalbC mice. We then demonstrated the capability of TrackUSF to detect and characterize, for the first time, modified social vocalization activity in *Shank3*-deficient rats, a rat model of ASD. We believe that this methodology would enable large-scale analysis of modified social vocalization activity in animal models of pathological conditions.

## Acknowledgments

This study was supported by The Human Frontier Science Program (HFSP grant RGP0019/2015), the Israel Science Foundation (ISF grants #1350/12, 1361/17), by the Milgrom Foundation and by the Ministry of Science, Technology and Space of Israel (Grant #3-12068).

## Materials and Methods

### Animals

#### Mice

BalbC and C57B animals were bred in a clean plastic chambers (GM500, Tecniplast, Italy) at 22°C and a 12-h light/12-h dark cycle (light on at 7 am) and received food and water ad libitum. All cages contain standard wood chip bedding, and cotton wool bedding material.

#### Rats

Subjects were naive Sprague Dawley (SD) male rats (8–12 weeks), commercially obtained (Envigo, Israel) and housed in groups of three to five animals per cage. *Shank3*-deficient rats, were a generous gift by Dr, Joseph Buxbaum at the Icahn School of Medicine at Mount Sinai. They were bred in a local colony and housed under the same condition described above. Wildtype (WT), heterozygous (Het) and knock-out (KO) littermates were offspring of heterozygous mating pairs. All rats were kept on a 12-h light/12-h dark cycle, light on at 9 p.m., with ad libitum access to food and water. Behavioral experiments took place during the dark phase under dim red light.

All experiments were approved by the Institutional Animal Care and Use Committee (IACUC) of the University of Haifa.

### Experiments

#### Mice

Vocal communications were recorded using a 1/4 inch microphone, connected to a preamplifier and an amplifier (Bruel & Kjaer) in a custom-built sound-shielded box. Vocalizations were sampled at 250 kHz with a CED Micro 1401-3 (Cambridge Electronic Design Limited, Sunnyvale, CA).

In each session, a pair of mice were kept in their home cage and the cage was placed within a custom soundproof box to minimize background noise. The microphone, inserted through the soundproof box lid, was suspended just above the home cage.

The system was programmed to record 10 minutes every hour for a duration of 12 hours that started after the first encounter between the animals. The audio clips were then analyzed offline in two stages (see below). Collectively, we recorded from 6 mice pairs. For each of these pairs, one clip that comprised the highest amount of vocalizations among all other clips was used for the comparison with the data obtained by our TrackUSF software.

#### Rats

The experimental setup consisted of a black Plexiglass arena (50 × 50 × 40 cm) placed in the middle of an acoustic chamber (90 × 60 × 85 cm). A compute-connected high-quality monochromatic camera (Flea3 USB3, Point Grey), equipped with a wide-angle lens, was placed at the top of the acoustic chamber, enabling a clear view and recording of the rat’s behavior using a commercial software (FlyCapture2, Point Grey). Video recordings were carried at a rate of 30 frames per second. Ultrasonic vocalizations were recorded using a condenser ultrasound microphone (Polaroid/CMPA, Avisoft) placed high enough above the experimental arena (Supplemental Fig. 2) so the receiving angle of the microphone can cover the whole arena. The microphone was connected to an ultra-sound recording interface (UltraSoundGate 116Hme, Avisoft), which was plugged into a computer equipped with the recording software Avi-soft Recorder USG (sampling frequency: 250 kHz; FFT-length: 1024points; 16-bit format). Synchronization between the video and audio recordings was achieved by making a hand click under the camera immediately after introducing the two rats to the arena.

### Video analysis (rats)

To track the interaction between dyads of same-age and same-genotype rats, we developed a new algorithm written in MATLAB (2017a) and added it to our previously published (Netser et al., 2017) Tracking software (“TrackRodent”, https://github.com/shainetser/TrackRodent). The main goal of this algorithm was to track the boundaries of the animals and determine for each frame whether they are merged (thus only one is detected = “interaction”) or not (thus two boundaries are detected = “no interaction”, Fig 6A).

All video analyses were done after correcting the raw behavioral data by skipping any gap of <?15 frames (0.5 s) in investigation and not considering it as breaking the investigation bout.

### Audio analysis

#### Manual analysis of mouse mating calls

Audio clips were analyzed offline in two stages. In the first stage single calls were extracted and the second stage of analysis was set to determine the spectral and temporal characteristics of each call in order to evaluate the vocal repertoire of the recorded mice. Spectral and temporal analysis of each mouse strain vocalizations was performed by creating spectrograms of the recorded files and marking the onset and offset of each calls within each recording. It was found in previous researches (Liu, Miller et al. 2003) that the minimal time interval between two distinct calls is approximately 800 msec, therefore we determined that a call ends and a new begins when there is at least an inter syllable interval of 1000 msec between the last syllable of the call and the first syllable of the following call. Time stamps were extracted and single call clips were created. These clips underwent further analysis of syllable extraction process in which every onset and offset of each syllable were listed to extract data regarding the temporal properties of the calls and aid in filtering the audio files and extracting spectral properties.

Filtering the audio vectors was possible using the syllable extraction process mentioned above, it allowed us to separate the audio files into two vectors: one containing emitted vocalizations with background noise and the second containing the inter-syllable intervals segments which contained only the background noise.

Subtracting the later vector from the former produced a cleaner audio vector that was used for spectral analysis. All vectors of ultrasonic vocalizations were normalized to the peak value in the range of 30-100 kHz.

### TrackUSF

#### Algorithm

Mel-frequency features represent the short-term power spectrum of a sound based on a linear cosine transform of a log power spectrum on a nonlinear Mel-scale of frequency. In this method, the frequency bands are equally spaced according to the Mel-scale. In our study, we expanded the Mel-frequency Cepstrum approach to represent rodents ultrasonic vocalizations using the Matlab function ‘mfcc’ (https://www.mathworks.com/matlabcentral/fileexchange/32849-htk-mfcc-matlab). We changed the lower and upper frequency limits to be between 15 kHz and 100 kHz, with the number of cepstral coefficients enlarged respectively. To analyze these features, we used dimensionality reduction by employing t-Distributed Stochastic Neighbor Embedding (t-SNE) algorithm (van der Maaten and Hinton, 2008), an algorithm that is particularly efficient for the visualization of high dimensional datasets. 3D t-SNE models each high dimensional object by a point in a three dimensions space to such a degree that similar objects are modeled by nearby points, while dissimilar objects are modeled by distant points with high probability. In our analysis, we used the function ‘tsne’ in Matlab with the default algorithm ‘barneshut’ and perplexity of 500 points.

The algorithm was embedded into a Graphical User Interface (GUI) written in MATLAB (Supplemental. Fig. 1 and the supplemental document “TrackUSF user manual”). Through the GUI, the user chooses sets of audio files for analysis (WAV format), divided according to their group identity. The analysis is then run and outputs the MFCC analyzed data, the t-SNE analyzed data, and the audio files in a MATLAB format (“.mat”). Next, the user can open those files for visualization of the data in 3D and for manually defining clusters on top of the t-SNE image result. After defining the clusters, the user can present the detected USFs on the spectrograms of the original data.

All variations of the software are deposited in GitHub under the following links: https://github.com/shainetser/TrackUSF

Spectrograms were computed using the standard ‘spectrogram’ function with a window of 512 samples, 50% overlap and sample rate of 250kHz. Power spectral density (PSD) for the different clusters was performed using a short-time Fourier transform with the same parameters as for the spectrograms. First, PSDs were performed for each USF separately. Then, the mean PSD for each cluster was calculated by averaging the PSDs of all USFs from the same cluster.

Calculating the probability of USFs occurrence relative to USFs from their own cluster or from other clusters was done in a time window of 6 msec for 0.5 sec before and after each USF detection. The synchronization between USFs and physical interaction was examined in a time window of 50 ms.

## Statistics

Data is presented as the mean ± SEM unless otherwise noted. Differences in the means of three or more groups were tested using one-way or two-way analysis of variance (ANOVA) followed by Bonferroni *post hoc* tests, when a significant main effect was found. In case of violation of ANOVA models assumptions (including lack of normal distributions), Kruskal-Wallis test was performed for comparing distributions of the groups, followed by a *post hoc* Dunn test with Bonferroni adjustment, when a significant result was found. When using a cutoff based on biological assumption, a Chi square test was performed. When significant results were obtained, additional Chi square tests were performed between each per of groups, adjusted by Bonferroni correction.

## Supplemental Figures

**Supplemental Figure 1:**
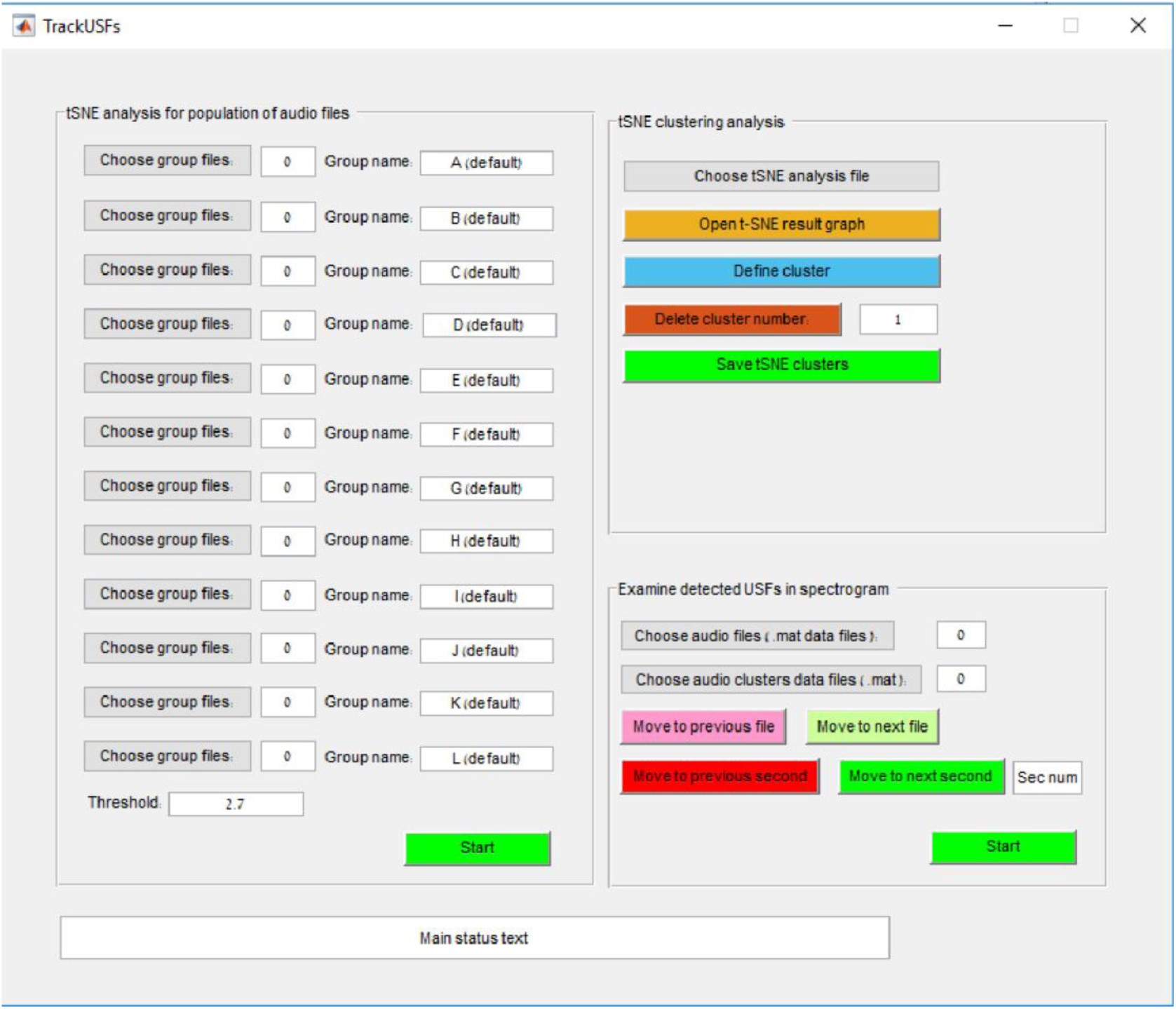
The Graphical User Interface (GUI) of the MATLAB-based TrackUSF software.

**Supplemental Figure 2:**
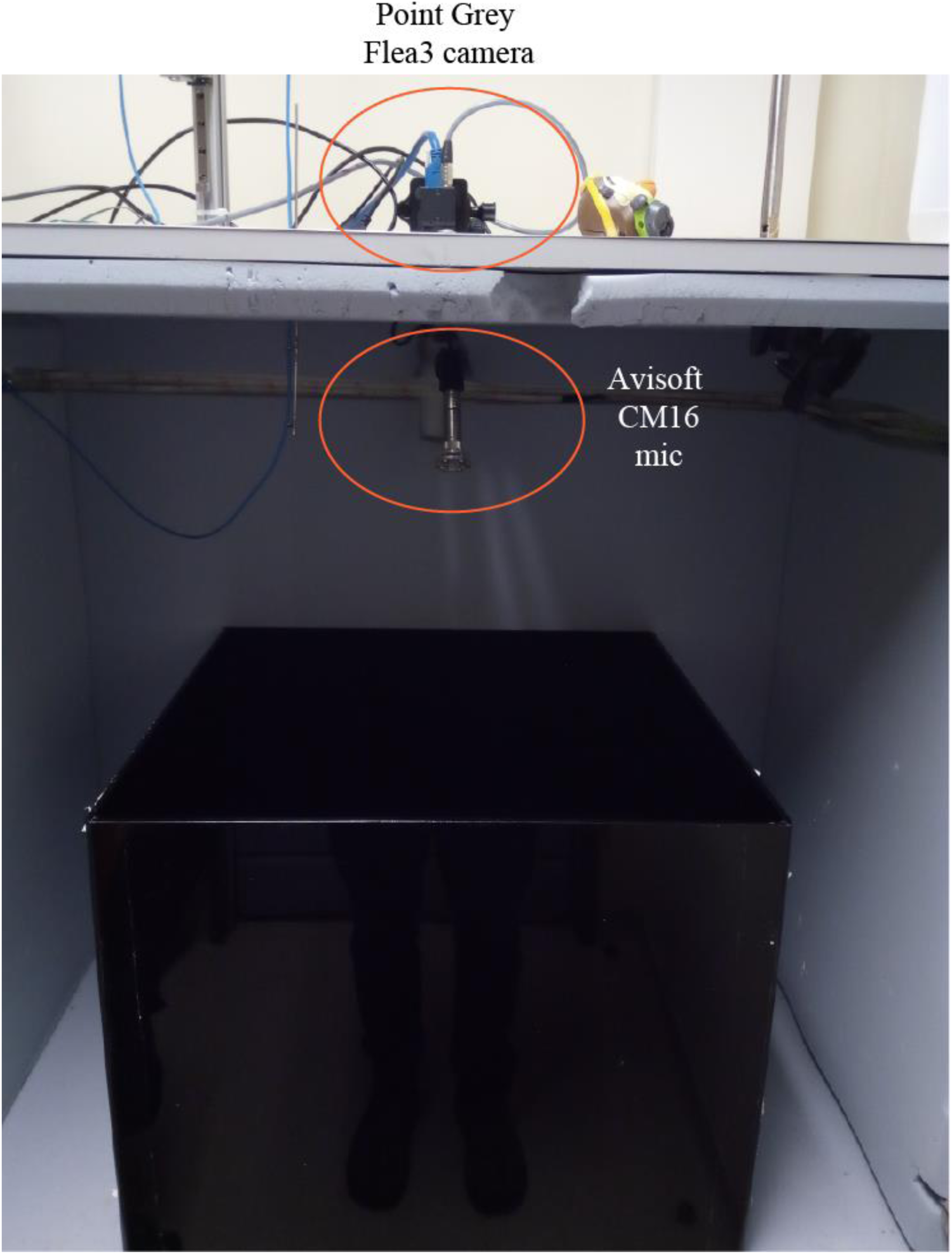
A picture depicting the experimental system used for recording social vocalizations in rats.

**Supplemental Figure 3:**
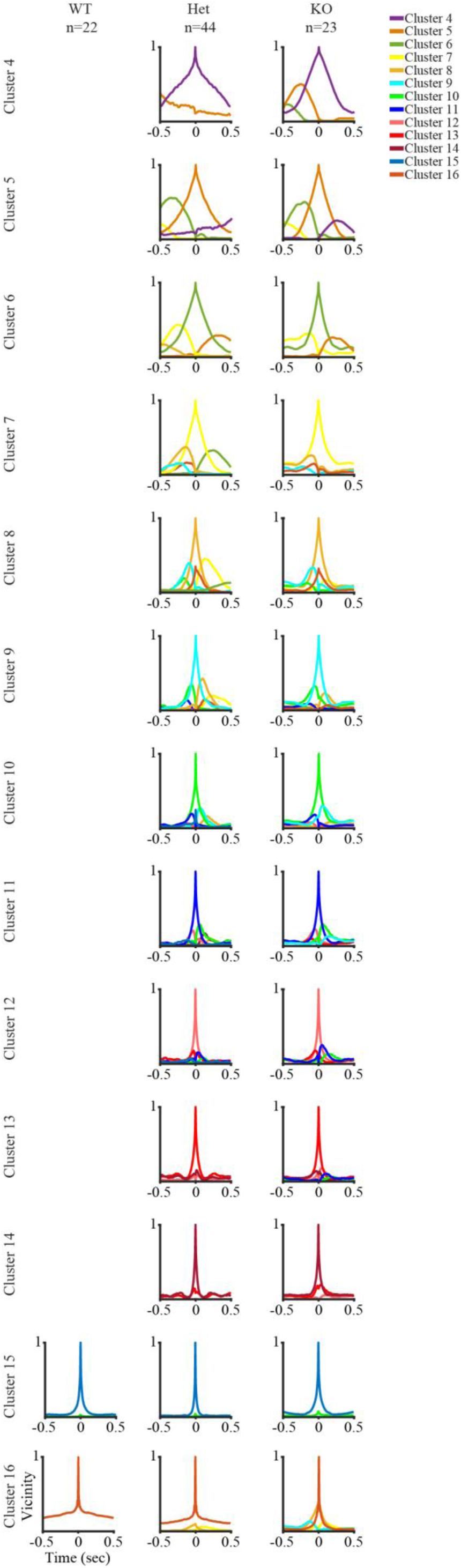
Vicinity curves for all clusters, across the three genotypes (WT-upper, Het-middle and KO-lower) of *Shank3*-deficient rats. In the case of WT animals, clusters 4-14 did not have enough representation for such analysis.

**Supplemental Figure 4:**
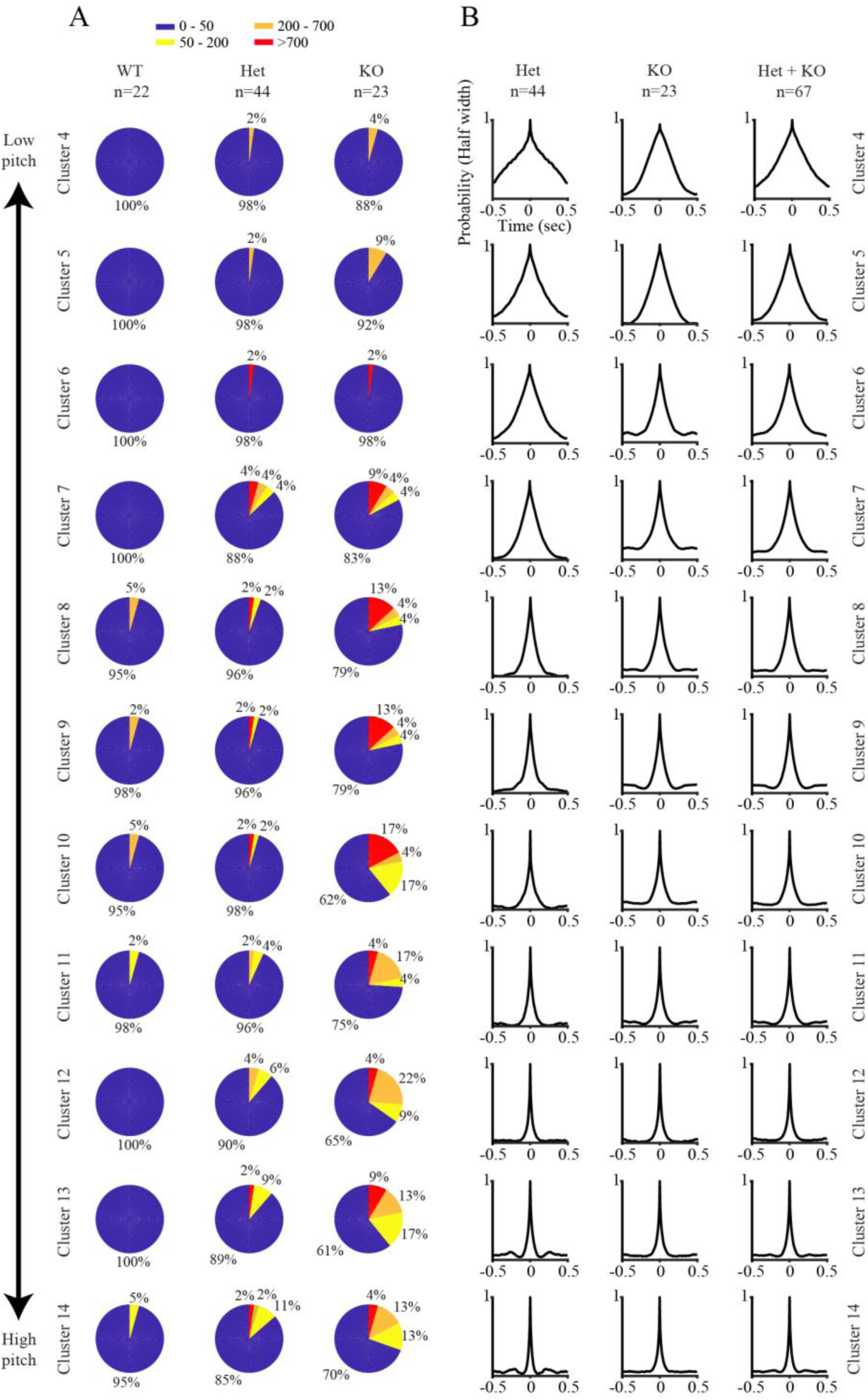
A) Proportions of sessions according to the numbers of USFs of clusters 4-14 they comprise, for the three genotypes of Shank3-deficient rats (novel animals and cagemates combined). B) Repeatability curves of clusters 4-14, for Het (left), KO (middle) and combined Het and KO animals (right). In the case of WT animals, clusters 4-14 did not have enough representation for such analysis.

**Supplemental Figure 5:**
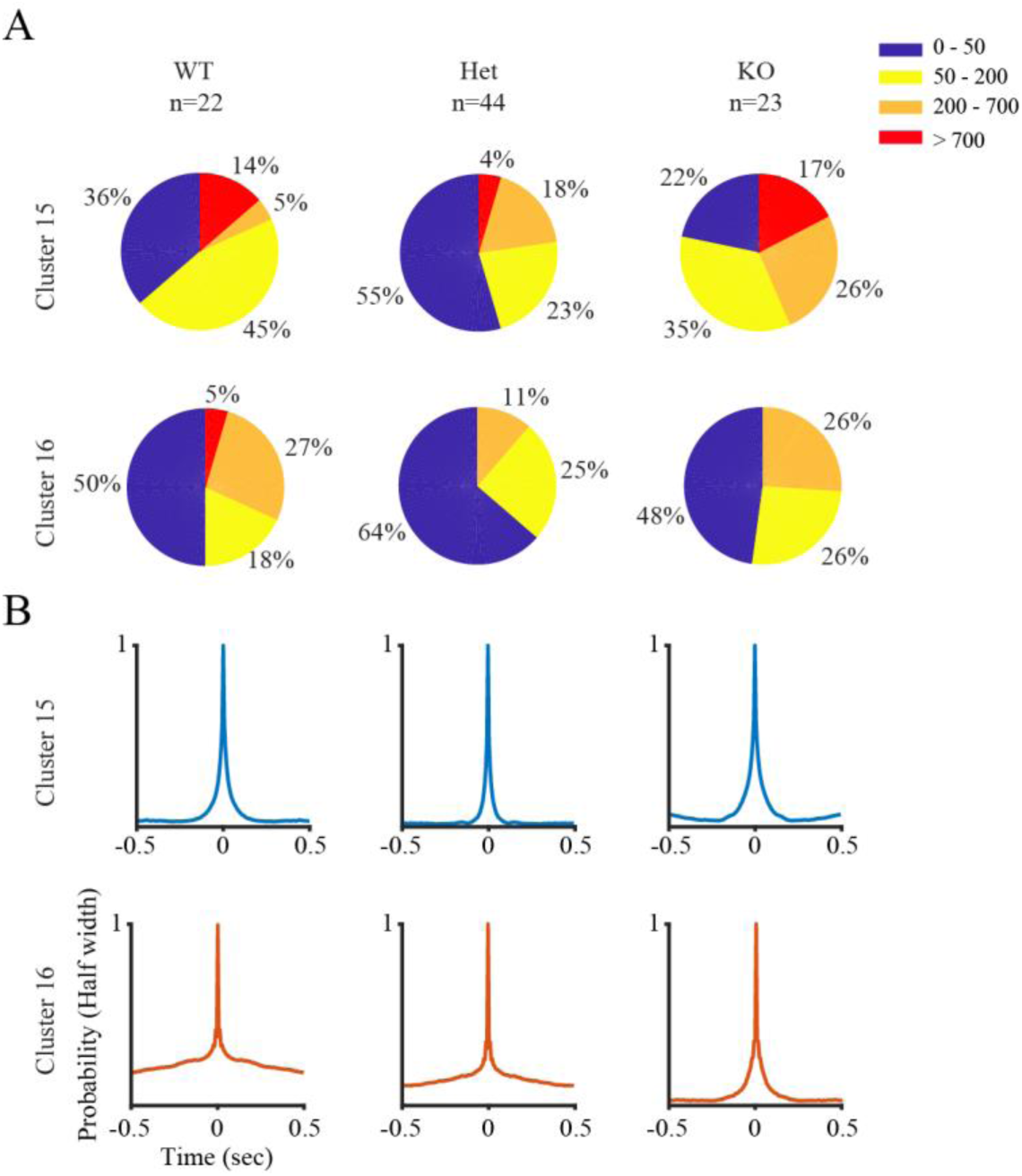
Proportions of sessions according to the numbers of USFs (A) and repeatability (B) of clusters 15-16 for all three genotypes of *Shank3*-deficient rats (see Fig. 5A-B for similar analyses of clusters 4-14).

